# Identification of a post-transcriptional regulatory element that responds to glucose in the African trypanosome

**DOI:** 10.1101/327346

**Authors:** Yijian Qiu, Vijay Shankar, Rooksana E. Noorai, Nelson Yeung, Sarah Grace McAlpine, James Morris

## Abstract

The ability to adapt to varying nutrient availability in changing environments is critical for successful parasitism. The lifecycle stages of the African trypanosome, *Trypanosoma brucei*, that infect the host mammalian bloodstream utilize glucose exclusively for ATP production. The finding that trypanosomes also inhabit other tissues that frequently contain lower glucose concentrations suggests blood stage parasites may have to respond to a dynamic environment with changing nutrient availability in order to survive. However, little is known about how the parasites coordinate gene expression with nutrient availability. Through transcriptome analysis, we have found blood stage parasites deprived of glucose alter gene expression in a pattern similar to transcriptome changes triggered by other stresses. A surprisingly low concentration of glucose (<10 μM) was required to initiate the response. To further understand the dynamic regulation of gene expression that occurs in response to altered glucose availability in the environment, we have interrogated the 3’UTR of cytochrome c oxidase subunit VI, a known lifecycle stage regulated gene, and have identified a stem-loop structure that confers glucose-responsive regulation at the translational level.

## INTRODUCTION

*T. brucei* encounters dramatic environmental changes and undergoes extensive physiological and metabolic alterations as it transits through its lifecycle in both the mammalian host and the tsetse fly vector. In the mammal, the bloodstream form (BF) parasites, which have been found to colonize a variety of tissues in addition to the bloodstream including the skin, brain, fat, and testis, respond to environmental cues with niche-specific responses (1-3). Similarly, in the fly vector, procyclic form (PF) trypanosomes sense their environment, modulating development into distinct tissue-dependent lifecycle stages in response to environmental cues (4).

Trypanosomes experience different glucose concentrations in the tissues of their mammalian host and tsetse fly vector, suggesting that the hexose abundance could serve as a cue to orient parasites to the identity of their environment. While BF parasites in the blood are exposed to glucose levels maintained at ∼5 mM, *T. brucei* can infect organs where there are lower glucose concentrations, including the brain (∼1 mM) and testes (0.1 mM) (5-8). Furthermore, parasites in the skin and fat tissue may encounter reduced glucose levels, which may have consequences for the parasites. For example, trypanosomes found in these tissues have distinct gene expression patterns that suggest they are able to metabolize alternative carbon sources (2). The health of the host may also influence glucose availability, as trypanosome infection can lead to changes in both blood and cerebrospinal fluid glucose levels (9-11). *In vitro* assays support the responsiveness of BF trypanosomes to altered glucose availability. For example, inhibition of glycolysis with the hexose transport inhibitor phloretin or 2-deoxy-glucose (2-DOG) has been shown to alter BF gene expression (12).

The gene expression changes in response to altered glucose availability are likely the consequence of post-transcriptional regulation, as *T. brucei* minimally regulates genes at the transcription level (13). This is due, in part, to the polycistronic nature of the organization of the parasite genome, which has multiple protein-coding genes clustered together that are co-transcribed by RNA polymerase II into polycistronic primary transcripts. These primary transcripts are then matured through the regulated processes of trans-splicing and polyadenylation into functional mRNAs (14). Mature transcripts are also subject to multiple levels of regulation, including alternative splicing, regulated transcript transport and subcellular localization, and degradation (15).

Key cellular components that participate in post-transcriptional regulation include RNA binding proteins, metabolites, and non-coding RNAs (16). In *T. brucei*, RNA binding proteins have been found to have a major influence on cell development and metabolism, including ZFP family members, ZC3H18, ALBA family members, RBP10, RBP6, DHH1, and hnRNP F/H (17-27).

The role of RNA binding proteins in influencing gene expression in response to environmental nutrient availability remains largely unresolved. Additionally, only a two *cis*-acting regulatory elements have been identified in *T. brucei* that are the basis for responses to specific environmental signals. The first, a 25-nucleotide 3’UTR element, is responsible for glycerol-induced changes in GPEET procyclin expression (28). In the second, a 35-nucleotide stem-loop in the 3’UTR of the NT8 purine transporter gene is responsible for repressed expression in response to guanosine (16). Other *cis*acting elements have been identified that are related to lifecycle stage-dependent expression. These include a 16-nucleotide element shared by several procyclin mRNAs (29), a 26-nucleotide stem-loop in the 3’UTR of *EP1* procyclin (30,31), a 7-nucleotide motif in PUF9 target mRNAs (32), and a 34-nucleotide element in the 3’UTR of ESAG9 family members transcripts (33).

Here, we describe the impact of changes to environmental glucose on BF parasites steady state transcript abundance. Additionally, we have identified a novel regulatory element in the 3’UTR of the cytochrome oxidase subunit VI (*COXVI*). This element, which is predicted to be a component of a stem-loop structure in the gene, regulates translation in response to glucose abundance.

## MATERIAL AND METHODS

### Parasite culture and treatment

Bloodstream-form (BF) 9013 parasites, a derivative of Lister strain 427 (34), were cultured in HMI9 (35) or glucose-deficient media (RPMIθ, ∼5 μM glucose) developed from (Hirumi et al. 1977) that includes glucose-free RPMI buffered to pH 7.4 with HEPES as a replacement for IMDM, elimination of SerumPlus, and use of dialyzed FBS (10% f.c., Corning). For treatments designed to remove glucose, parasites in log phase were collected (800 x g, 8 min) and washed three times in PBS before being resuspended at 2-6 × 10^5^ cell/mL in pre-warmed RPMIθ supplemented with or without 5 mM glucose.

### RNA analysis

RNA was isolated from frozen treated cells that were collected by centrifugation (800 x g, 8 min), washed in PBS, before being flash frozen and stored at -80°C. Total RNA was isolated using an Aurum Total RNA Mini Kit (Bio-Rad) and DNA was removed by incubation with DNase I (RT, 15 min) in one of the kit steps. RNA samples were quantified using an Eon Microplate Spectrophotometer (BioTek) to ensure all samples had an A260/A280 value of ∼2. Real-time quantitative PCR was performed using a Verso 1-step RT-qPCR Kit (ThermoFisher) using the protocol provided by the manufacturer in a CFX96 Touch™ Real-Time PCR Detection System (Bio-Rad). Primers were designed using the GenScript Real-time PCR (TaqMan) Primer Design (https://www.genscript.com/tools/real-time-pcr-taqman-primer-design-tool) and blasted against the trypanosome genome at TritrypDB (http://tritrypdb.org/tritrypdb/) to ensure specificity. RNA (100 to 500 pg) and 300 nM of each primer were added to a 10 µL reaction. The reaction protocol was as follows: cDNA synthesis (50°C, 10 min); reverse transcripate inactivation (95°C, 5 min); PCR and detection (10 sec, 95°C and 30 sec, 60°C, for 45 cycles; melt curve analysis (1 min, 95°C, 1 min, 55°C, and 10 sec from 55 to 95°C for 80 cycles with a 0.5°C increment each cycle). Only products with single and clearly defined melting curves were used for analysis. The 60S ribosomal protein L10a (RPL10A) or telomerase reverse transcriptase (TERT) genes were used as expression references to solve transcript Ct values using the comparative Ct (2^-ΔΔCT^) method (Schmittgen and Livak 2008; Brenndorfer and Boshart 2010).

For RNAseq, libraries were constructed using the TruSeq Stranded mRNA Library Prep Kit (Illumina, San Diego, CA USA) and input RNA normalized to 200 ng per sample. Sequencing was performed on either an Illumina HiSeq2500 (Hollings Cancer Center, Medical University of South Carolina) at 2 × 125 paired-end reads or an Illuimina NextSeq 550 (Genomics & Computational Laboratory, Clemson University) at 2 x150 paired-end reads. FastQC (http://www.bioinformatics.babraham.ac.uk/projects/fastqc) and Trimmomatic (36) were used to assess quality metrics with low quality bases and any adapter sequences removed. GSNAP (37) was used to index and align reads to the *T. brucei* TREU927 reference genome minus the 11 bin scaffold/chromosome (http://tritrypdb.org). Subread’s featureCounts (38) was used to count uniquely mapped, properly paired reads per gene for each sample. The Bioconductor package edgeR (39,40) was used to calculate differential gene expression and produce plots.

Hierarchical clustering of the log_2_ fold change in gene expression induced by glucose depletion in BF and LS, and comparison of blood to fat conditions in LS (2) was performed in Genesis (41) using the average linkage WPGMA clustering algorithm. To construct Venn diagrams, gene lists from the differential gene expression analysis across all three comparisons were filtered for FDR-adjusted p < 0.05, then separated into upregulated and downregulated sets and further partitioned into subsets based on their presences in comparisons and combinations of comparisons.

### Constructs and luciferase assays

A trypanosome luciferase expression vector was generated by cloning the luciferase open reading frame into pXU, a pXS6-derived plasmid modified by elimination of the trypanosome UTRs flanking the multi-cloning site (Supplementary Figure 1). The 5’UTR sequences were cloned between the ClaI and BamHI sites or introduced by PCR, while different 3’UTRs were cloned into the XhoI and HindIII sites. Plasmids were linearized by NotI and transfected into BF as described (42).

Luciferase activity was measured using a slightly modified protocol from (16). Briefly, 2 × 10^6^ cells were pelleted (800 x g, 8 min, RT) before being washed once in PBS. Cell pellets were resuspended in 100 μL of lysis buffer (100 mM potassium phosphate, pH 7.8, 2 mM dithiothreitol (DTT), 0.1% Triton X-100, 10% glycerol.), vortexed, and incubated on ice (5 min). Debris was removed by centrifugation (16,000 × g, 10 min, 4°C) and the supernatant was transferred to pre-chilled tubes. Luciferase assays were then performed using the Luciferase Assay System (Promega) in a BioTek™ Synergy™ H1 Hybrid plate reader according to the manufacturer’s direction and relative luciferase activity normalized to cell number.

## RESULTS

### Assessment of BF viability in the absence of glucose

Glucose is a critical carbon source for BF trypanosomes. To understand how BF parasites respond to glucose depletion, the viability of cells grown in media with different concentrations of the hexose was determined. When BF parasites were seeded at 3 × 10^5^ cells/mL in very low glucose media (RPMIθ, ∼ 5 μM glucose) supplemented with 5 mM glucose, viability was maintained and cell division was detectable after six hours (Figure 1A). If supplemental glucose was not added, BF parasites did not proliferate but most (∼80%) remained viable for at least 12 hours. After 12 hours, both cell number and the percentage of viable cells decreased (Figure 1B).

**Figure 1.**
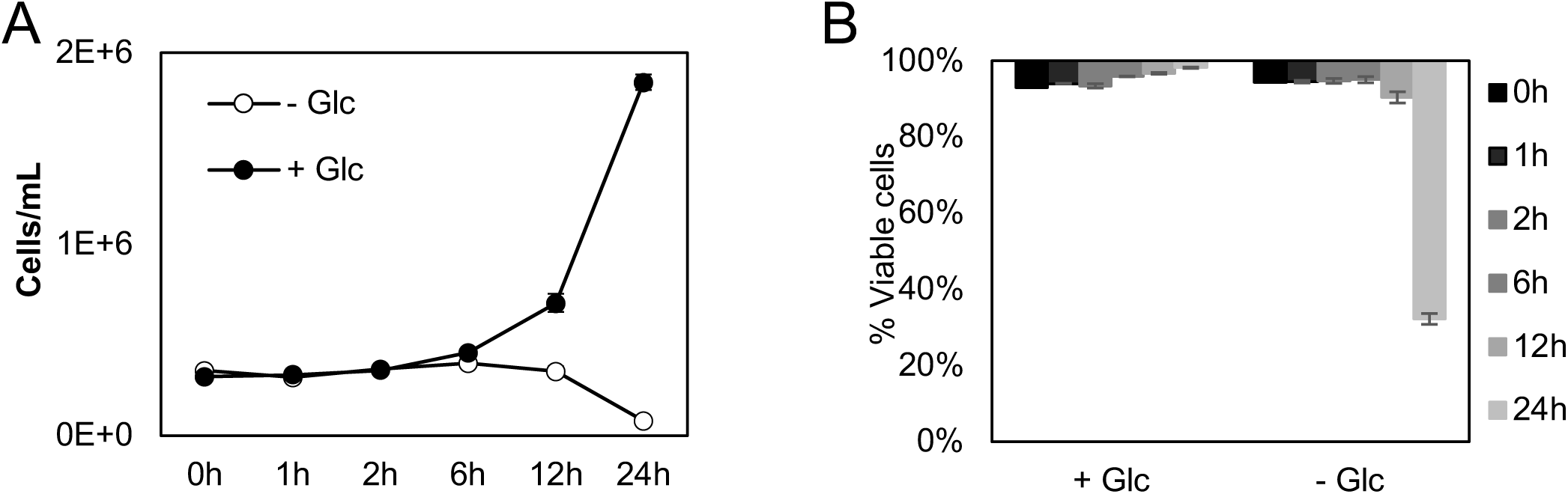
BF cells are viable in the near-absence of glucose for at least 12 hours. (A) Growth curve of BF *T. brucei* cells in RPMIθ supplemented with or without 5 mM glucose. Cells were seeded at 3 × 10^5^/mL and counted at the indicated times. Bars indicate standard deviation. (B) Assessment of the viability of cells during the treatment by propidium iodide staining. Glc, glucose.

### Transcriptome analysis of bloodstream form trypanosomes in glucose-rich and deficient conditions

The period of time, when BF parasites viability was largely unaffected by the near-absence of glucose, offered an opportunity to score parasite response to the treatment. To analyze this response, high-throughput sequencing (RNAseq) was performed on BF incubated for 12 hours in very low glucose media supplemented with 5 mM or 1 mM glucose. While there were almost no differences between the transcriptomes of cells grown in 5 mM and 1 mM glucose, cells cultured in the near absence of glucose (∼5 μM) had 1882 transcripts that were differentially expressed with at least a 2-fold change in expression level (and FDR < 0.05). Of the differentially regulated genes, 1282 were upregulated and 600 were downregulated (see Supplementary Table 1 for the full list).

Differentially expressed transcripts were mapped to Kyoto Encyclopedia of Genes and Genomes (KEGG) pathways (43) through TritrypDB (44) and analyzed for Gene Ontology (GO) enrichment. When the KEGG groups were ranked based on the significance of fold-enrichment, transcripts of TCA cycle-associated genes were among the most significantly upregulated while genes associated with glycolysis, an indispensable pathway for BF parasite ATP generation, were among the most down-regulated (Figure 2 and Supplementary Table 2). These changes in the expression of metabolic genes were similar to those observed in BF treated with phloretin (12). GO enrichment analysis revealed a significant upregulation of transcripts encoding transporters on cell surface, as well as those involved in regulation of macromolecule metabolic process and gene expression at posttranscriptional level (Figure 3A). Genes encoding glycosomal components as well as proteins related to unfolded protein binding, RNA binding, and aminoacyl-tRNA ligase activity were downregulated (Figure 3B).

**Figure 2.**
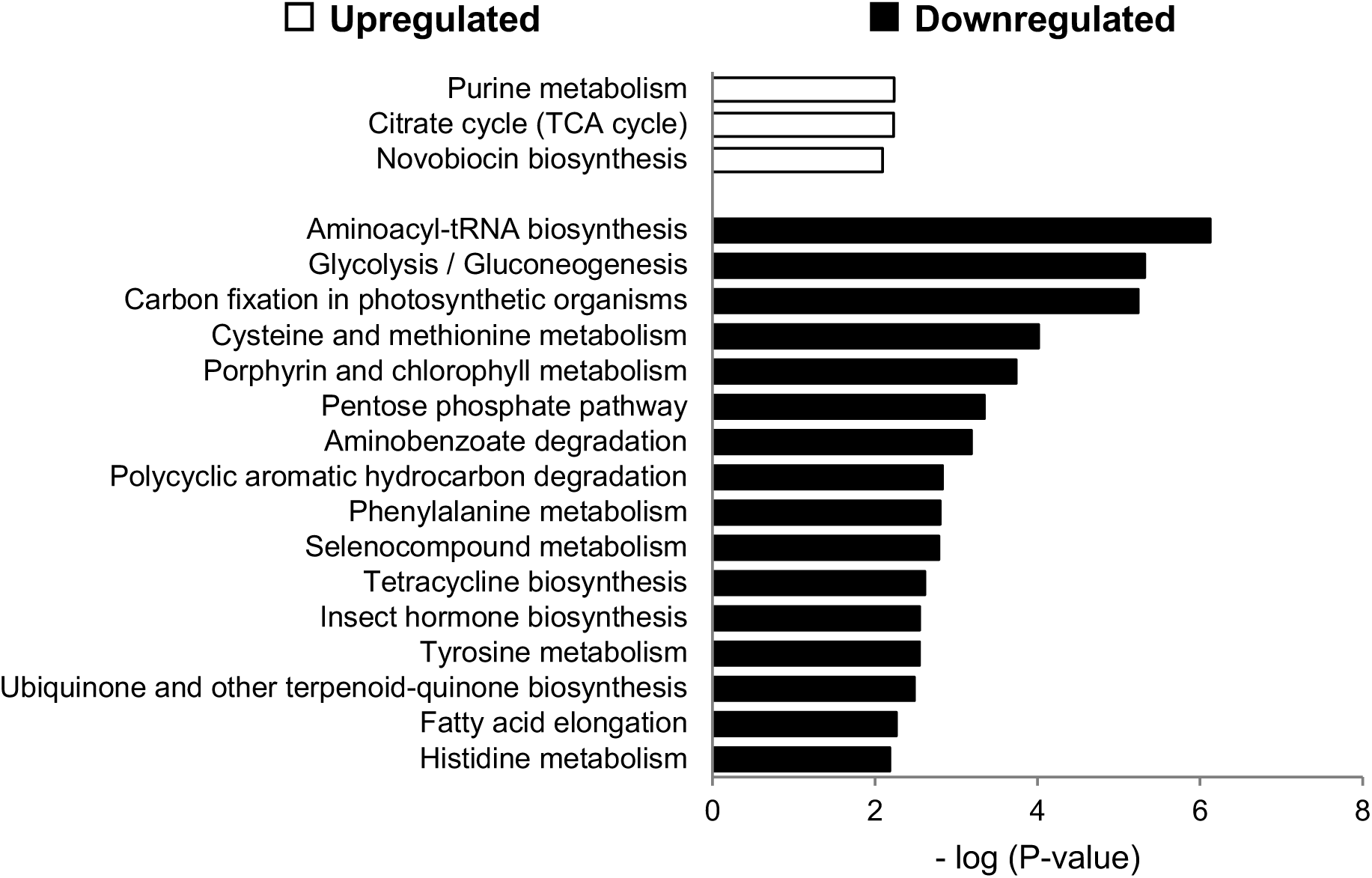
Transcriptome remodeling of metabolic pathways in BF parasites when glucose levels were reduced to ∼5 µM. Significantly regulated genes were mapped to the Kyoto Encyclopedia of Genes and Genomes (KEGG) pathways. Only genes that were overrepresented in up- and down-regulated pathways are shown (p-value < 0.01).

**Figure 3.**
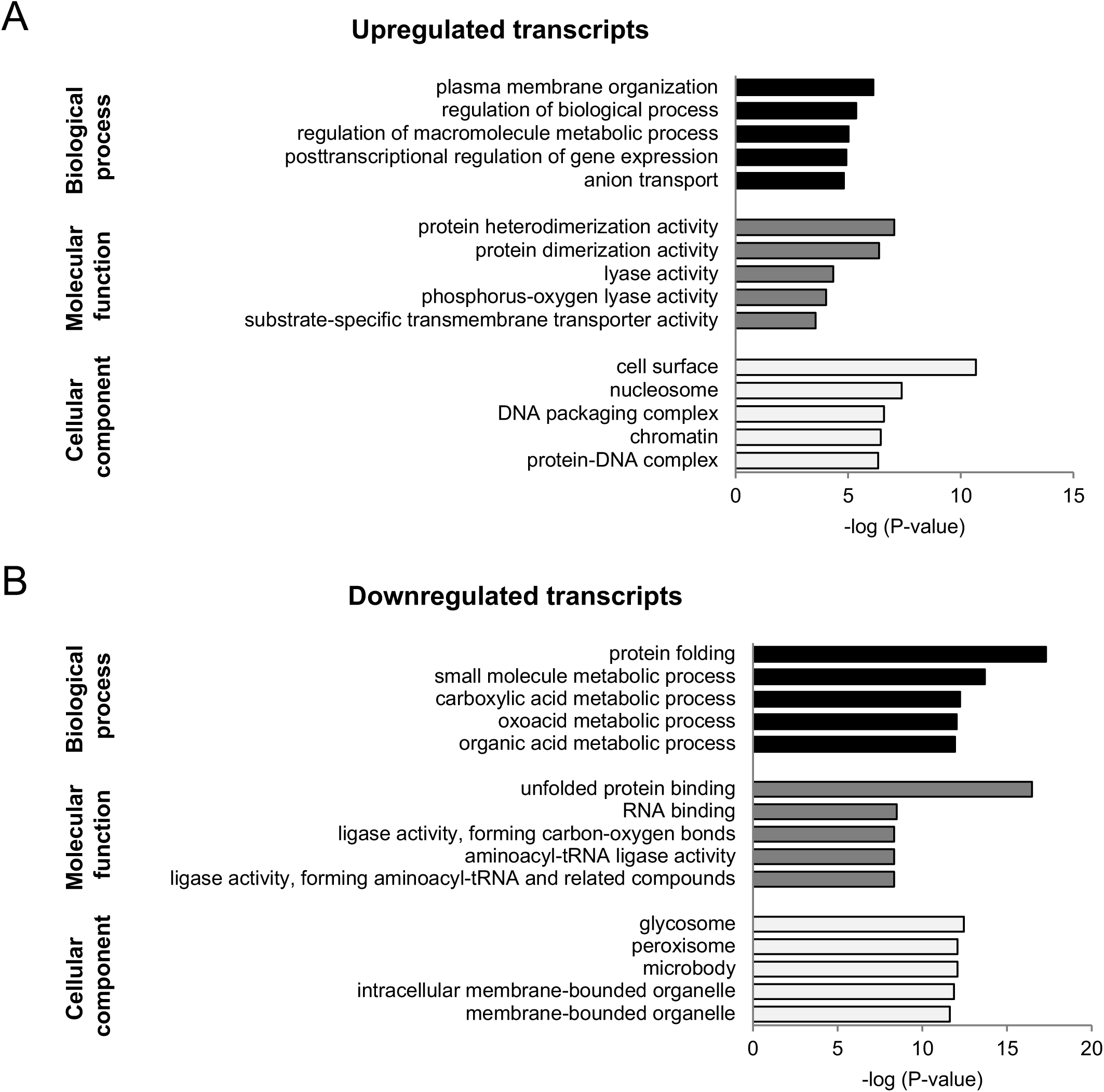
Gene Ontology (GO) enrichment of significantly regulated transcripts in BF parasites when glucose was removed. Only the top five GO non-general terms in each category are included (p-value < 0.001). For a complete list, please see Supplementary Table 3.

Notably, several developmentally-regulated genes were also found in the list of glucose-responsive differentially-expressed genes. These included several significantly upregulated transcripts encoding proteins essential for differentiation, like *PAD2* (Tb927.7.5940), *PAD1* (Tb927.7.5930) and *RBP6* (Tb927.3.2930), as well as developmentally-regulated metabolism genes that are involved in oxidative phosphorylation like *COXVI* (Tb927.10.280), *COXVII* (Tb927.3.1410), and *COXVIII* (Tb927.4.4620) (Supplementary Table 1). Glycosome-resident protein genes that function in aspects of glucose metabolism, another developmentally-regulated metabolic pathway, were significantly downregulated. These included the genes for enolase (Tb927.10.2890), fructose-bisphosphate aldolase (Tb927.10.5620) and glyceraldehyde 3-phosphate dehydrogenase (Tb927.6.4300).

Since glycolysis is critical to BF trypanosomes for ATP production, the observed transcriptome regulation after glucose removal could reflect a response to general cellular stress. To begin to address this possibility, BF cells were treated with blasticidin, an antibiotic that inhibits protein translation and represses cell growth. Cells were treated with the drug and harvested when the majority of the population was still viable (Supplementary Figure 2) and the expression pattern of select genes was assessed. By quantitative reverse transcriptase-PCR (qRT-PCR), most transcripts tested after either blasticidin treatment or culture in the near-absence of glucose were similarly regulated, suggesting the observed changes in these cases was not specific to the reduction of glucose. However, hexokinase 1 (*HK1*), trypanosome hexose transporter 1 (*THT1*), EP procyclin (*EP*), and GPEET procyclin (*GPEET*) were among the genes that were differentially regulated in cells cultured in the near-absence of glucose (Figure 4) but either not regulated or regulated in a distinct manner in response to blasticidin treatment.

**Figure 4.**
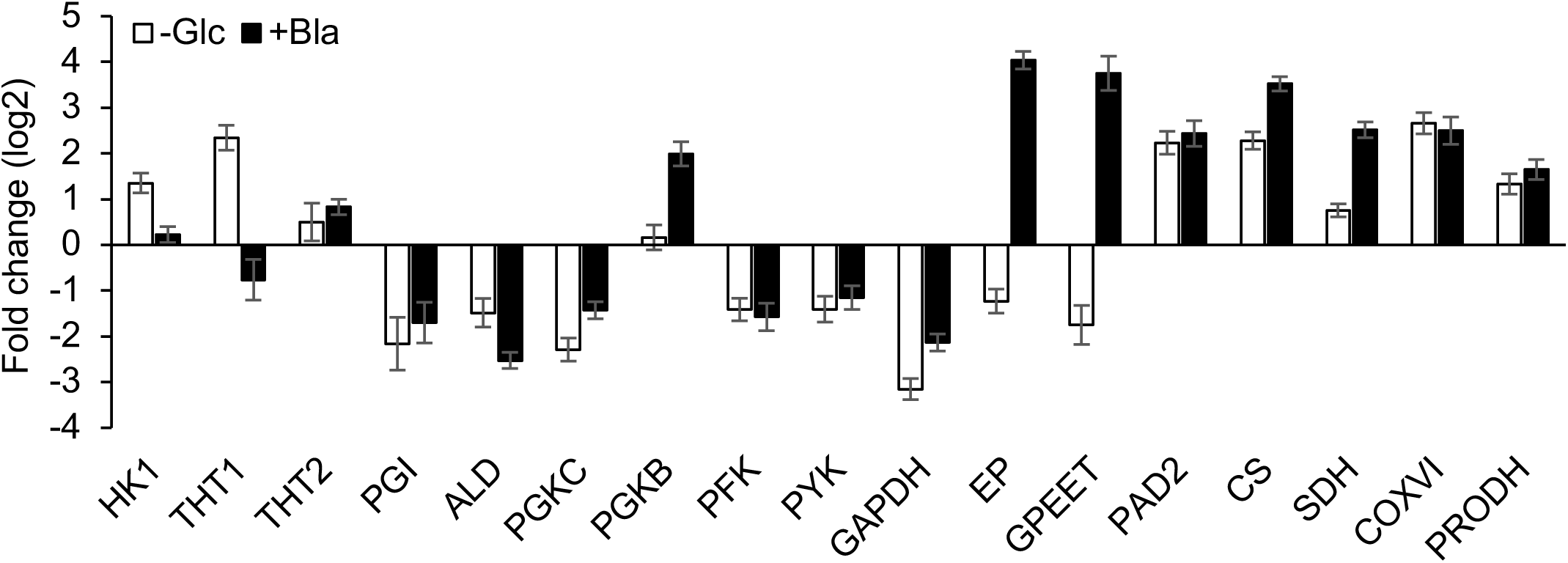
Comparison of the impact of glucose depletion to treatment with blasticidin on relative transcript abundance in BF parasites. Total RNA was isolated as described in the Material and Methods and analyzed by qRT-PCR. The expression levels of transcripts from biological triplicates were compared to their levels in untreated cells to generate fold-change values. Triangles indicate genes that are differentially regulated in response to glucose depletion when compared to those regulated in response to blasticidin treatment. Error bars indicate standard deviations in triplicate. - Glc, glucose depletion treatment; +Bla, blasticidin treatment; HK1, hexokinase 1; THT1, glucose transporter 1; THT2, glucose transporter 2; PGI, glucose-6-phosphate isomerase; ALD, fructose-bisphosphate aldolase; PGKC, phosphoglycerate kinase C; PGKB, phosphoglycerate kinase B; PFK, ATP-dependent phosphofructokinase; PYK, pyruvate kinase; GAPDH, glyceraldehyde 3-phosphate dehydrogenase; EP, EP procyclin; GPEET, GPEET procyclin; PAD2, protein associated with differentiation 2; CS, citrate synthase; SDH, succinate dehydrogenase; COXVI, cytochrome oxidase subunit VI; PRODH, proline dehydrogenase.

### Glucose-responsive regulation is rapid and requires an unexpectedly low level of glucose

To study the timing and glucose concentration required for regulation induced by glucose depletion, three significantly upregulated transcripts, *HK1, COXVI*, and *RNA binding protein 5* (*RBP5)* were selected as reporter genes. The transcript abundance of these reporter genes was scored from BF cells incubated in RPMIθ for one, two, six, and twelve hours and compared to those seeded in RPMIθ with 5 mM of glucose. Transcripts of all three genes were found to be upregulated one hour after treatment (Figure 5A), suggesting the regulation is rapid. To assess the glucose concentration required for the observed regulation, BF cells were incubated in media with different amounts of glucose (5, 10, 55, and 500 µM) for two hours followed by assessment of reporter transcript abundance. Surprisingly, significant up-regulation of the reporter genes was not observed unless the glucose concentrations were below 10 μM (Figure 5B).

**Figure 5.**
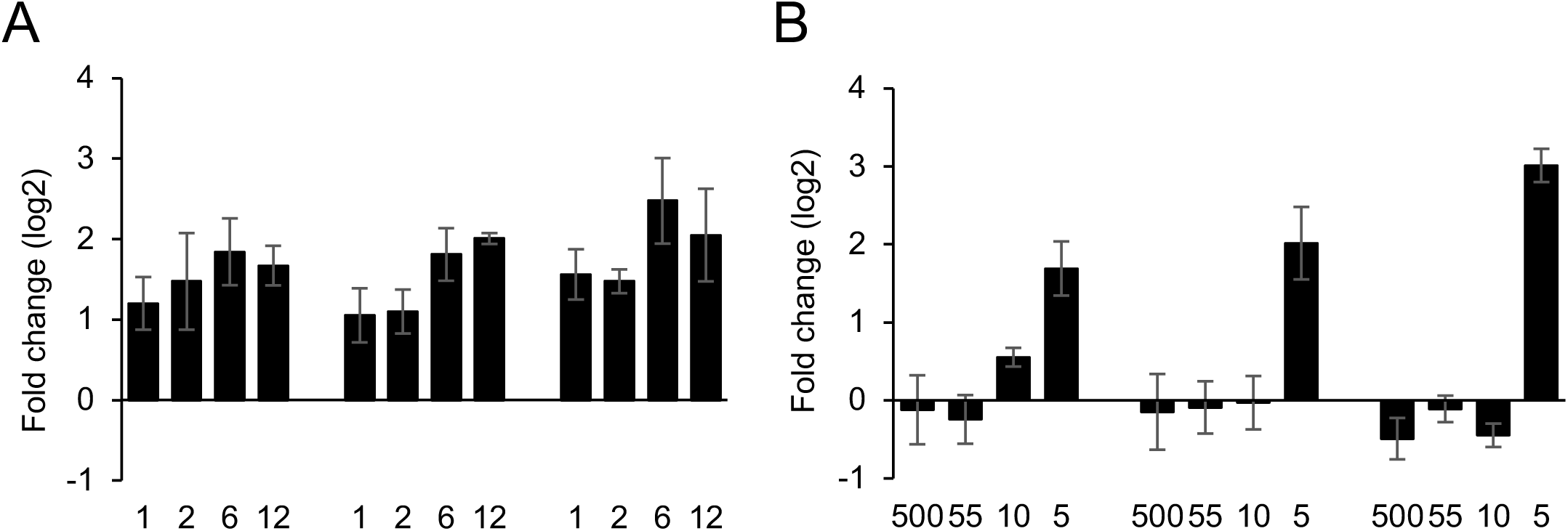
The glucose-related regulation requires an extremely low level of glucose. The expression changes of three genes known to be upregulated in response to glucose depletion were analyzed by qRT-PCR. The expression level of each transcript was compared to its level in untreated cells to generate a fold-change. (A) The regulation of the three genes was scored in BF cells maintained in glucose-depleted media (f.c. of glucose ∼5 µM) for 1, 2, 6, and 12 hours. (B) The expression of the three reporter genes in BF cells incubated in RPMIθ media contains 500 μM, 55 μM, 10 μM, and 5 μM of glucose, respectively. Error bars indicate standard deviations in triplicate.

### The 3’UTR of COXVI is responsible for glucose-dependent regulation

Transcript steady-state abundance in Kinetoplastida parasites is primarily controlled by post-transcriptional means and the responsible regulatory genetic elements are frequently located in the 3’UTR of the regulated genes (45). Cytochrome oxidase subunit VI (*COXVI)* encodes a product that is essential for the electron transport chain, the expression of which is largely repressed in BF (46). The 3’UTR of *COXVI* has been studied in the context of lifecycle stage-specific regulation, with regulatory regions identified that primarily operate at the translational level (31). To determine if the 3’UTR of *COXVI* harbors similar elements responsible for the observed upregulation after glucose depletion, the sequence was cloned downstream of a luciferase gene reporter and stably integrated into BF cells. The 5’UTR of *COXVI* was also tested using the same system, with the 5’UTR of the EP1 procyclin gene and 3’UTR of the actin gene used as controls.

Cells harboring stably-integrated luciferase constructs were cultured in glucose-replete conditions and then washed, resuspended in RPMIθ or RPMIθ supplemented with 5 mM glucose and incubated for two hours prior to RNA analysis (Figure 6A) or testing for luciferase activity (Figure 6B). Surprisingly, luciferase steady state transcript levels were not influenced by the presence of either the *COXVI* 5’ or 3’UTR (Figure 6A), which suggested that the observed changes in steady state levels of *COXVI* were the result of a regulatory element within the open reading frame of the gene.

**Figure 6.**
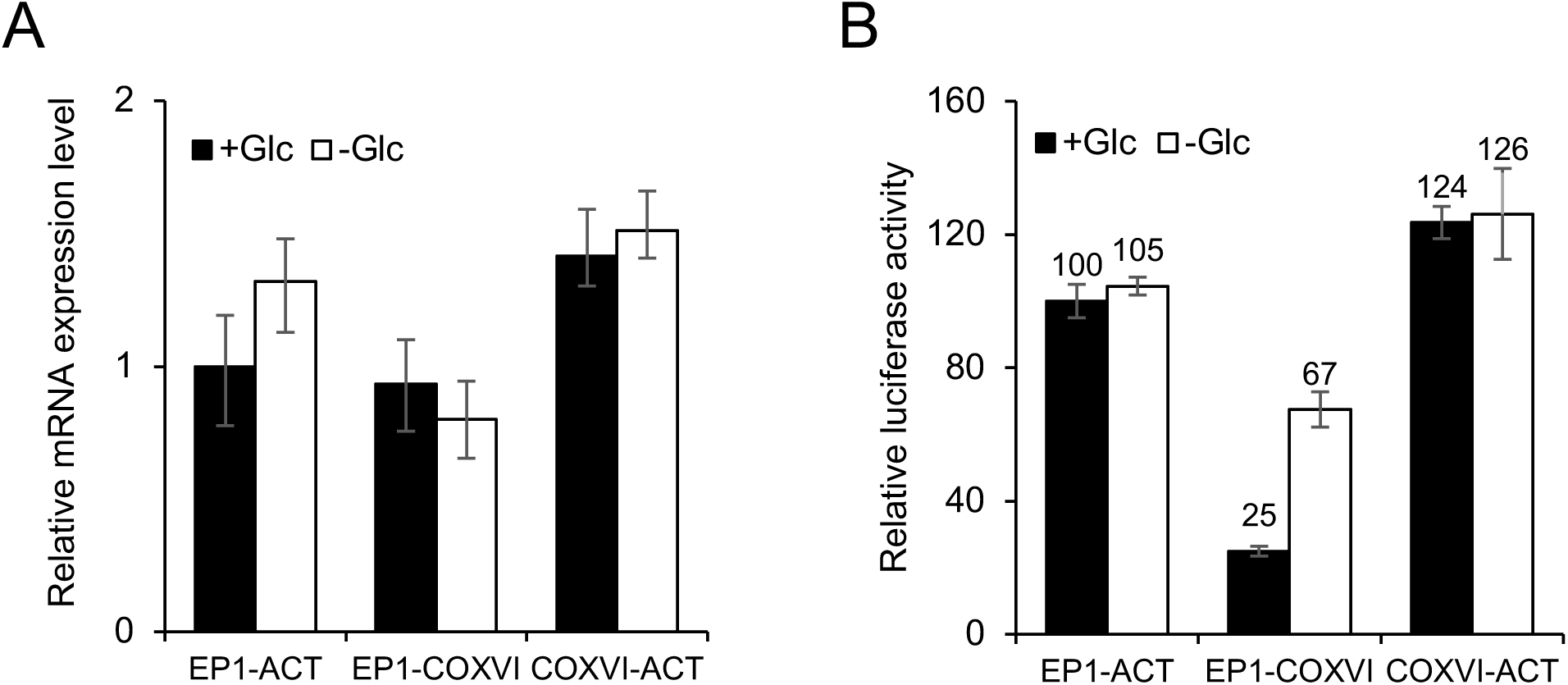
The *COXVI* 3’UTR regulates luciferase activity at the protein level. BF cells where stably transformed with pXU bearing the luciferase ORF cloned between the indicated 5’ and 3’UTRs from *EP1, actin a* (*ACT*) and *COXVI*. The *EP1* 5’UTR and *actin a* 3’UTR were used as controls. Steady-state transcript abundance (A) and luciferase activity (B) were assessed. In both cases, control cells in glucose-replete media were used as references for normalization. Bars represent standard deviation in triplicate. ** indicates at least 2-fold change in activity with p-value < 0.01 according to two-sided student t-test.

Nevertheless, luciferase activity was increased almost three-fold from constructs harboring the *COXVI* 3’UTR luciferase, indicating the potential presence of an element that influences protein expression in the 3’UTR of the gene (Figure 6B). Unlike the 3’UTR, the *COXVI* 5’UTR had no impact on differential expression in response to glucose.

### Identification of a regulatory element within the COXVI 3’ UTR

The length (187 nucleotides) and relatively simple predicted secondary structure of the *COXVI* 3’UTR served as the starting point to identify the glucose responsive regulatory element that was responsible for the increased luciferase expression. Using the RNAstructure web server (http://rna.urmc.rochester.edu/RNAstructureWeb/Servers/Predict1/Predict1.html) (47), the *COXVI* 3’UTR structure was modeled, which resulted in a hairpin-like structure with the first and last ∼60 nucleotides of the UTR partially annealing to form the stem. The intervening region from nucleotides 77-124 was predicted to form two short stem-loops (SLI and SLII, Figure 7A).

**Figure 7.**
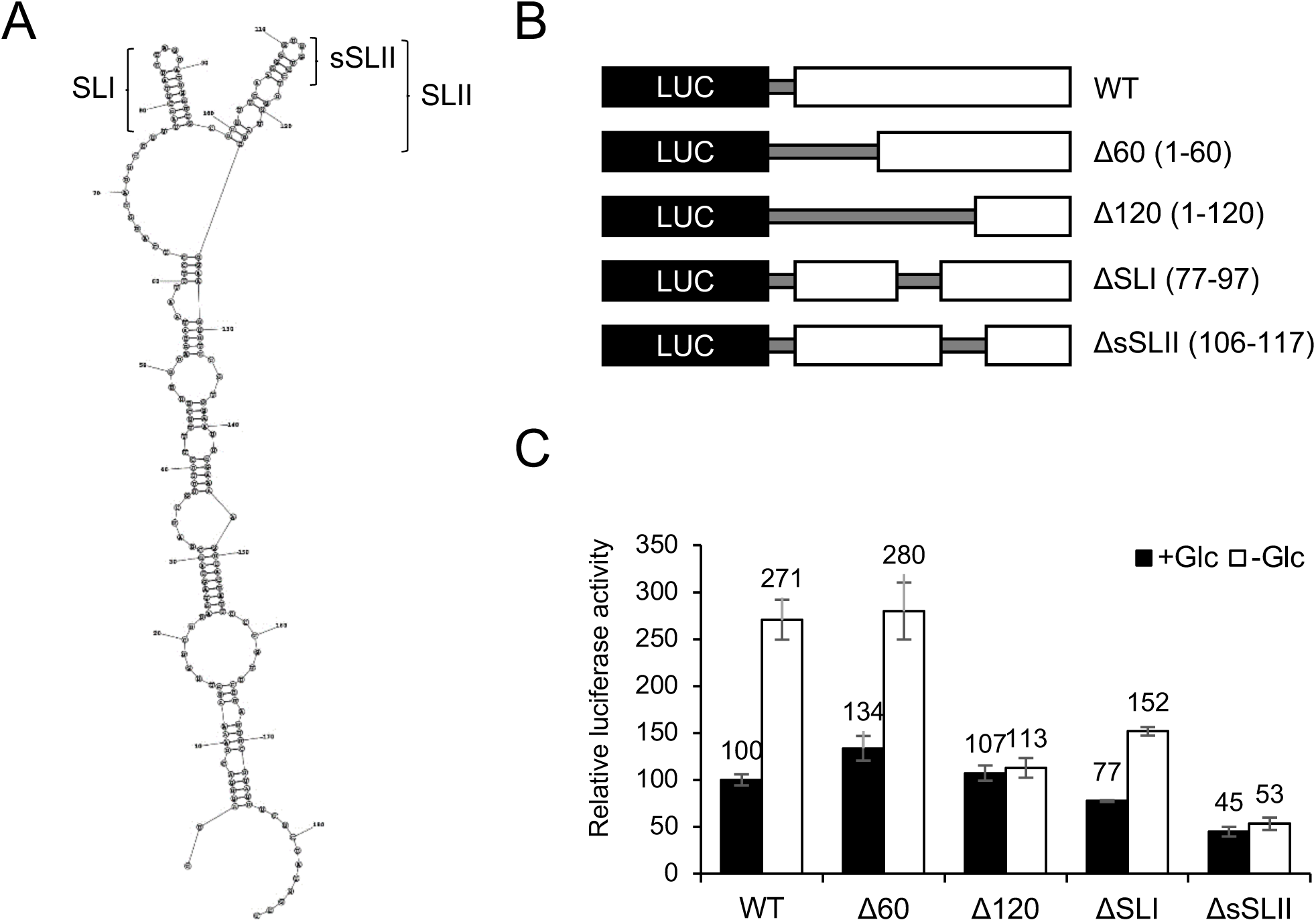
Resolving components of the *COXVI* 3’UTR that impart regulation in response to glucose depletion. (A) The predicted maximum free energy secondary structure of the *COXVI* 3’UTR. Stem-loop I (SLI) and stem-loop II (SLII) are indicated, with an additional feature at the apex of SLII was noted as sSLII. (B) Schematic of the constructs used to explore the role of the *COXVI* 3’UTR on expression. The *EP1* 5’UTR was used in all constructs and deletions (Δ) and position of the eliminated nucleotides are indicated. (C) Luciferase activity was measured for all variants. Relative activity levels were normalized to signal from cells cultured in glucose-rich (5 mM) media expressing luciferase fused to the full-length *COXVI* 3’UTR. Bars represent standard deviation. ** indicates at least 2-fold change in activity with p-value < 0.01 according to two-sided student t-test.

To test the importance of these putative structures in the glucose-responsive increase in expression, modified *COXVI* 3’UTRs were generated by mutation and fused to the luciferase gene (Figure 7B). Modeling of these altered 3’UTRs suggests the changes had a variable impact on the overall structure (Supplementary Figure 3). Removal of the first 60 nucleotides of the 3’UTR (Δ60) did not alter the glucose-responsive expression of luciferase activity. However, differential expression was abolished after removal of the first 120 nucleotides (Δ120, Figure 7C). This result suggested the region between nucleotides 60 and 120, which includes the stem-loops SLI and SLII in the structural model, contains a potential regulatory element.

### SLII mediates changes in gene expression in response to glucose-deficiency

To study whether SLI or SLII between nucleotides 77-124 were necessary for glucose-responsiveness, constructs bearing luciferase under the regulation of a 3’UTR missing either stem-loop were generated by mutagenesis and tested for their impact on mRNA steady state levels and luciferase expression.

In the absence of the first stem-loop (ΔSLI), the glucose-responsive increase in luciferase activity was not altered (Figure 7C). However, elimination of 12 nucleotides from the apex of SLII, to yield ΔsSLII, did abolish the response of the element. This 12-mer was insufficient to impart glucose-dependent regulation to 3’UTRs not normally subject to such regulation. For example, the addition of sSLII to that of actin a (*ACT*) or the 60S ribosomal protein L10a (*RPL10A*) 3’UTRs did not alter luciferase expression glucose-depleted conditions (Figure 8A). However, modification of the *ACT* or *RPL10A* 3’UTRs by addition of the 26-nucleotide SLII was sufficient to impart glucose-sensitive regulation to luciferase expression (Figure 8A.)

**Figure 8.**
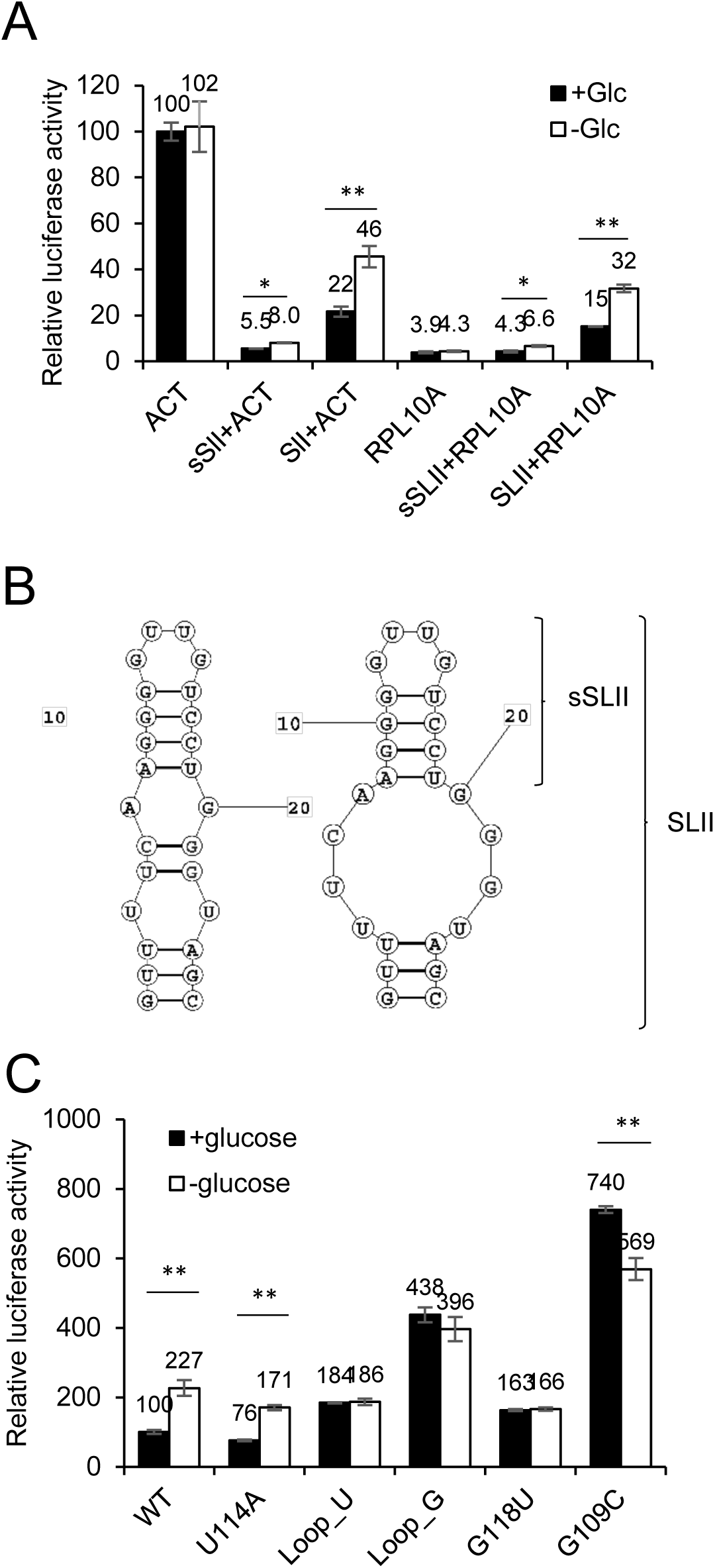
Identification of structure and sequence features in SLII that are required for responsiveness to glucose depletion. (A) Luciferase activity measured from trypanosomes expressing luciferase under the regulation of the *actin a* (ACT) or *60S rRNA L10A* (RPL10A) 3’UTRs with or without sSLII or SLII appended at the 5’ end of the 3’UTR. (B) Prediction of two possible secondary structures of SLII. The apex of SLII, coined sSLII, is indicated. (C) Luciferase activity from cells expressing luciferase under the regulation of mutant *COXVI* 3’UTRs. Relative activity levels were normalized to the activity scored from parasites expressing luciferase fused to the *actin a* 3’UTR or wild type *COXVI* 3’UTR in glucose-replete media. Bars represent standard deviation in triplicate. ** indicates at least 2-fold change in activity with p-value < 0.01 according to two-sided student t-test. * indicates less-than-2-fold change in activity but with p-value < 0.01.

To resolve the nucleotides in SLII that were essential for the glucose responsiveness of the element, mutations were made in the SLII in the context of full-length *COXVI* 3’UTR and luciferase expression assessed. (For a full list of secondary structure predictions for these variant UTRs, see Supplementary Figure 4). Mutation of uracil 114 in the *COXVI* 3’UTR into adenine increased the number of nucleotides in the loop and had little impact on the upregulation of luciferase (Figure 8C). All other mutations, including replacement of the two guanines on the loop with uracils (resulting in a loop composed of only uracils), replacement of the two uracils on the loop by guanines (to yield a loop of only guanines), mutation of G118 into uracil (changing the bubble adjacent to the loop into a stem), and mutation of G109 into cytosine (which disrupted the whole SLII structure), abolished the upregulation of luciferase in cells deprived of glucose (Figure 8C). These results suggested the four nucleotides on the loop, the adenine or guanine in the bubble, and the overall structure were important for the regulation.

## DISCUSSION

The mammalian infectious stage of the African trypanosome occupies diverse niches that have different levels of available glucose. While the average glucose level in blood plasma is maintained at ∼5 mM, other tissues contain a more dynamic range of this hexose. For example, brain glucose is only ∼10-20% of the levels found in blood (7,48), while the glucose concentration in the seminiferous tubular fluid of the testes is even lower (less than 2% that of blood) (3,8,49). BF parasites incubated in the near-absence of glucose remained viable for half a day during which time the expression of many lifecycle stage transcripts were altered, a behavior usually noted as the parasite transits to the next host, the tsetse fly.

The differentiation-like trend in transcriptome change was also observed in BF parasites treated with phloretin (12), although there were some differences in transcript profiles between the treatments including the changes in the expression of *EP, GPEET, THT1*, and *HK1*. Stage-specific transcripts were also differentially regulated in response to other treatments. Treatment with the antibiotic blasticidin (Figure 4), which interferes with translation, or 2-DOG, a competitive inhibitor of hexokinases (12) (Supplementary Figure 5) altered the expression of developmentally regulated transcripts. Similarly, manipulation of inositol polyphosphate multikinase (IMPK) levels in BF parasites has recently been shown to trigger a series of gene expression changes consistent with progress toward differentiation to PF parasites, such as the loss of VSG and expression of EP procyclin (50). Together, these observations suggest an overall common response among BF trypanosomes that encounter different potentially life-threatening stresses that includes altered expression of developmentally regulated genes.

The response to glucose depletion may be, in part, tailored to the low glucose conditions the parasite encounters *in vivo* in certain tissues. For example, the upregulation of THT1 and HK1 may be an effort to improve uptake and utilization of glucose as concentrations of the hexose fall, while the concurrent downregulation of procyclin genes suggests the parasite pauses the developmental reprogramming required to differentiate to the next lifecycle stage in the nutrient-deficient environment. Because these studies used lab-adapted monomorphic BF parasites that are only partially able to respond to authentic environment cues that are known to trigger pleomorphic parasites to differentiate (51,52), it is possible that these responses reflect unique behaviors of laboratory strains.

Comparison of the transcriptional profiles of monomorphic BF and long slender (LS) pleomorphic parasites reveals that the two lines respond with overall similar changes in gene expression after culture in glucose depleted conditions (Figure 9). This observation suggests that while monomorphic BF cells are impaired in their ability to differentiate, they retain the pathways necessary for response to environmental glucose. The pattern of expression in response to environmental glucose may be useful in identifying the nature of authentic cues in specific tissues. For example, pleomorphic LS parasites from adipose tissue likely do not use glucose as a cue for the regulation of gene expression in response to their environment. These cells, which upregulate genes involved in major metabolic pathways like glycolysis, amino acids metabolism, and fatty acid °-oxidation (2), share limited gene expression profile similarity with the profiles from BF or LS cultured in glucose-depleted conditions (Figure 9A and B).

**Figure 9.**
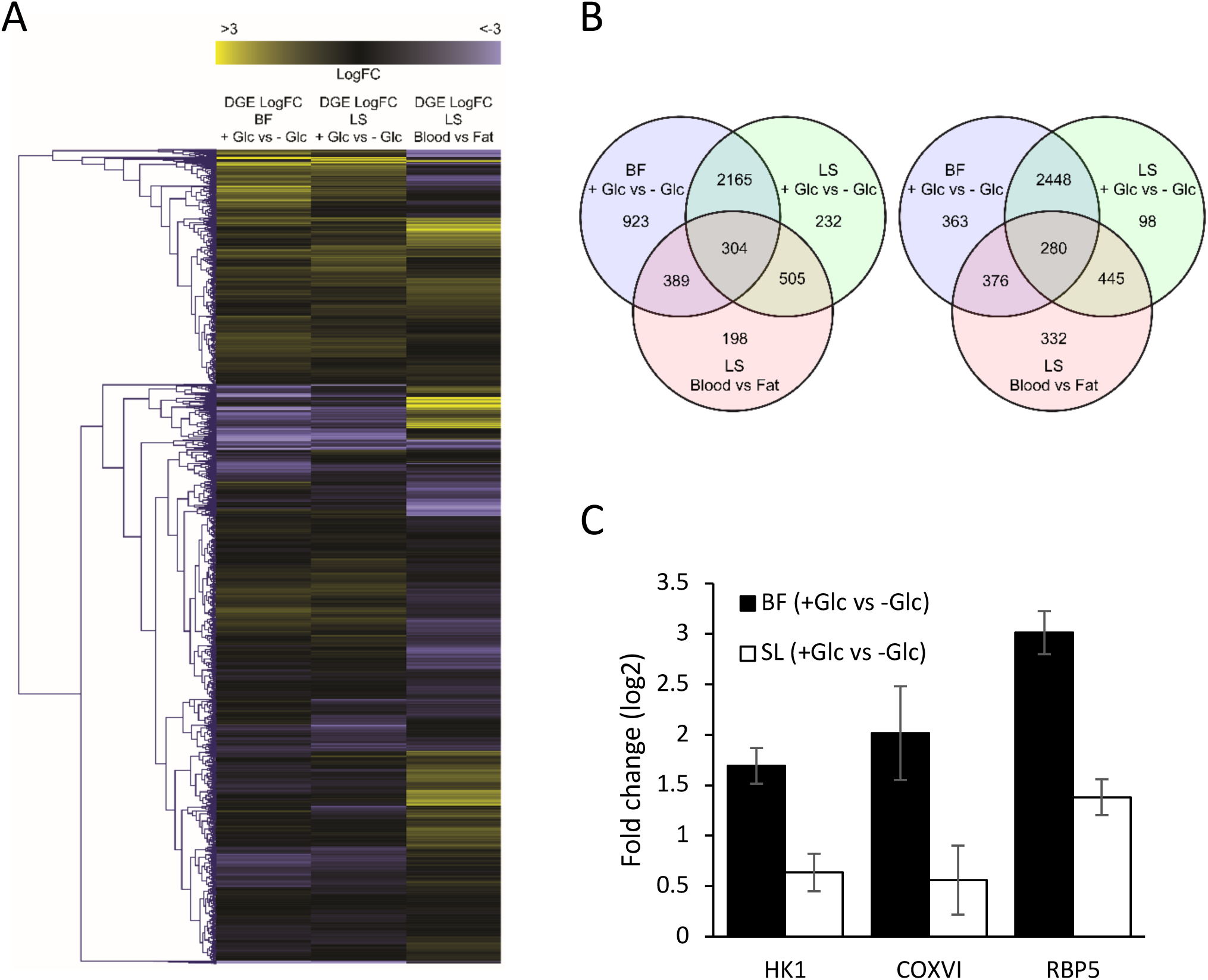
Comparative transcriptome analysis for blood stage parasites in different environments. (A) Hierarchically clustered heat map comparing the overall transcript regulation pattern among monomorphic BF and pleomorphic LS cells treated in glucose depletion media, and LS cells harvested from adipose tissue (2). The differentially expressed gene (DEG) fold-changes (FC) were obtained by comparing the gene expression levels of cells in a glucose depleted versus a glucose replete environment and in adipose tissue versus blood. (B) Venn diagrams of upregulated and downregulated genes in the three different cases above. (C) The regulations of the three genes in BF and LS cells maintained in glucose-depleted media were scored by qRT-PCR. The expression levels of transcripts were compared to their levels in untreated cells to generate fold-change values. Error bars indicate standard deviations in triplicate.

The regulation of transcript abundance in response to glucose depletion is rapid, taking place within hours of treatment, which is consistent with the observation that developmentally related transcripts tend to have short half-lives as a result of regulation of mRNA decay rates (53). Curiously, the glucose concentration required to observe the regulation was lower than 10 μM. Mammalian tissues tend to have higher available glucose concentrations, but parasites in tissues that already have extremely low glucose level (like the testes) may deplete glucose to levels low enough to initiate the response (46,54). Interestingly, this concentration is also below the *K*M values for all known glucose binding or transporting proteins in *T. brucei* (55-57). Two such 3’UTR elements have been identified, indicating the existence of an unknown mechanism of glucose perception in the parasite.

The alteration of translation in response to glucose depletion has been extensively studied in yeast (58-61) and found to occur extremely rapidly (within 5 minutes) upon change in carbon source availability (62). Glucose depletion similarly led to a rapid (∼20 minute) alteration of translation in a mammalian neural cell line (63), with TCA cycle and oxidative phosphorylation being among the most impacted pathways. These observations suggest that the oabserved altered translation of the *T. brucei* COXVI subunit may be the result of a potentially conserved regulatory mechanism.

In the glucose-poor tsetse fly tissues, mitochondrially-based metabolism is important for success of the parasite, which is reflected in the upregulation in PF parasites of components of amino acid metabolism pathways. For example, the cytochrome c oxidase complex is highly expressed in PF parasites and lacking in proliferative BF parasites (64). The 3’UTRs of COX family members have been studied in the context of life stage developmental regulation (31) and an 8-nucleotide element (UAUUUUUU) in genes of the family has been described. While *COXVI* transcripts harbor a shortened version of this element (UAUUUUU) in the last 60 nucleotides of the 3’UTR, this sequence is not responsible for the regulation induced by glucose removal. Deletion of the first 120 nucleotides of the *COXVI* 3’UTR, which preserved the UAUUUUU, eliminated the glucose responsiveness of the UTR (Figure 7C).

Another consensus sequence, UAG(G)UA(G/U), has also been identified in six members of the *COX* family (31). The *COXVI* 3’UTR contains two versions of this sequence. The first one, UAGUAG, is located at nucleotides 22–27 of the *COXVI* 3’UTR, while the second, UAGUAU, (nucleotides 86–91) makes up part of SLI. Deletion of the first was found to increase reporter protein activity in BF ∼3-fold (31). In this study, the deletion of the first 60 nucleotides (eliminating the first UAG(G)UA(G/U)), or deletion of SLI (which eliminates the second UAG(G)UA(G/U)), only slightly altered the steady-level abundance of luciferase (Figure 7C). Moreover, the glucose-depletion induced regulation was largely maintained in these two constructs (Figure 7C), indicating UAG(G)UA(G/U) was not responsible for the regulation triggered by glucose removal. Interestingly, the deletion of the first UAG(G)UA(G/U) introduced a new stem-loop structure into the 3’UTR based on secondary structure modeling (Supplementary Figure 3). The same stem-loop structure was not formed when either the first 60 nucleotides or the SLI of *COXVI* 3’UTR were removed (Supplementary Figure 3) and may have been important for the previously observed increase in reporter activity (31).

Structural features are believed to be central for the recognition of RNA regulatory elements (RREs) by RNA binding proteins (RBPs) (65). RBPs typically do not access primary nucleotide sequence information in regions of double-stranded structure but rather recognize sequences in single-stranded regions (66). The secondary structures of transcripts have been found to impact translational efficiency (67-70). In our study, single-stranded regions of the SLII of *COXVI* 3’UTR were important for sugar-based regulation, as mutations that altered either the single-strand loop or bubble of SLII impacted the response. Additionally, changing the structural context of the critical nucleotides in the SLII altered element function. This was demonstrated by the loss of regulation noted after engineering a single nucleotide mutation predicted to change the overall structure of the 3’UTR. The only variant that maintained normal glucose-responsive protein expression resulted in a predicted enlarged loop with unaltered single-stranded recognition sequences at the apex of the loop.

In this study we discovered a novel 26-nucleotide regulatory element in the *COXVI* 3’UTR that responds to the near-absence of glucose by increasing protein translation without a corresponding increase in steady-state transcript abundance. This regulatory element is characterized by an imperfect short stem with a loop at its apex containing the tetrameric nucleotide sequence GUUG. Although we have not identified sequences with high identity to the element in other transcripts, stem-loops with similar structural and sequence features have been identified in other upregulated transcripts like *HK1, PAD2, CS, COXVII* and *COXVIII* (Supplementary Figure 6). The role of these in translational regulation remains to be resolved.

Our findings suggest that African trypanosomes respond to general translational stress or external glucose depletion through a regulatory network that influences the stability and translation of certain groups of transcripts required for adaptation. Understanding the adaptive behavior of parasites under these stressful conditions, particularly during drug treatment, could reveal means for the development of targeted strategies to combat drug resistance, especially for those drugs that target critical metabolic pathways.

## AVAILABILITY

FastQC is available at http://www.bioinformatics.babraham.ac.uk/projects/fastqc. The RNAstructure web server is available at http://rna.urmc.rochester.edu/RNAstructureWeb/Servers/Predict1/Predict1.html. The *T. brucei* TREU927 reference genome is available at http://tritrypdb.org. GenScript Real-time PCR (TaqMan) Primer Design is available at https://www.genscript.com/tools/real-time-pcr-taqman-primer-design-tools.

### DATA AVAILABILITY

RNAseq data are available from NCBI under BioProject (PRJNA454021: *Trypanosoma brucei* Raw sequence reads (Taxid:5702)).

### SUPPLEMENTARY DATA

Supplementary Data is available as a separate PDF.

## ACKNOWLEDGEMENT

The authors would like to thank Dr. Chris Saski and Jillian Milanes for their technical assistance.

## References

1. Capewell, P., Cren-Travaille, C., Marchesi, F., Johnston, P., Clucas, C., Benson, R.A., Gorman, T.A., Calvo-Alvarez, E., Crouzols, A., Jouvion, G. et al. (2016) The skin is a significant but overlooked anatomical reservoir for vector-borne African trypanosomes. Elife, 5.

2. Trindade, S., Rijo-Ferreira, F., Carvalho, T., Pinto-Neves, D., Guegan, F., Aresta-Branco, F., Bento, F., Young, S.A., Pinto, A., Van Den Abbeele, J. et al. (2016) *Trypanosoma brucei* parasites occupy and functionally adapt to the adipose tissue in mice. Cell Host Microbe, 19, 837–848.

3. Claes, F., Vodnala, S.K., van Reet, N., Boucher, N., Lunden-Miguel, H., Baltz, T., Goddeeris, B.M., Buscher, P. and Rottenberg, M.E. (2009) Bioluminescent imaging of *Trypanosoma brucei* shows preferential testis dissemination which may hamper drug efficacy in sleeping sickness. Plos Neglect Trop D, 3.

4. Sharma, R., Gluenz, E., Peacock, L., Gibson, W., Gull, K. and Carrington, M. (2009) The heart of darkness: growth and form of *Trypanosoma brucei* in the tsetse fly. Trends Parasitol, 25, 517–524.

5. McNay, E.C. and Gold, P.E. (1999) Extracellular glucose concentrations in the rat hippocampus measured by zero-net-flux. J Neurochem, 72, 785–790.

6. Levin, B.E. (2000) Glucose-regulated dopamine release from substantia nigra neurons. Brain Res, 874, 158–164.

7. de Vries, M.G., Arseneau, L.M., Lawson, M.E. and Beverly, J.L. (2003) Extracellular glucose in rat ventromedial hypothalamus during acute and recurrent hypoglycemia. Diabetes, 52, 2767–2773.

8. Robinson, R. and Fritz, I.B. (1981) Metabolism of glucose by Sertoli cells in culture. Biol Reprod, 24, 1032–1041.

9. Moon, A.P., Williams, J.S. and Withersp.C. (1968) Serum biochemical changes in mice infected with *Trypanosoma rhodesiense* and *Trypanosoma duttoni*. Exp Parasitol, 22, 112-&.

10. Batista, J.S., Rodrigues, C.M., García, H.A., Bezerra, F.S., Olinda, R.G., Teixeira, M.M. and Soto-Blanco, B. (2011) Association of *Trypanosoma vivax* in extracellular sites with central nervous system lesions and changes in cerebrospinal fluid in experimentally infected goats. Vet Res, 42, 63.

11. Bisoffi, Z., Beltrame, A., Monteiro, G., Arzese, A., Marocco, S., Rorato, G., Anselmi, M. and Viale, P. (2005) African trypanosomiasis gambiense, Italy. Emerg Inf Dis, 11, 1745.

12. Haanstra, J.R., Kerkhoven, E.J., van Tuijl, A., Blits, M., Wurst, M., van Nuland, R., Albert, M.A., Michels, P.A.M., Bouwman, J., Clayton, C. et al. (2011) A domino effect in drug action: from metabolic assault towards parasite differentiation. Mol Microbiol, 79, 94–108.

13. Schwede, A., Kramer, S. and Carrington, M. (2012) How do trypanosomes change gene expression in response to the environment? Protoplasma, 249, 223–238.

14. Preusser, C., Jae, N. and Bindereif, A. (2012) mRNA splicing in trypanosomes. Int J Med Microbiol, 302, 221–224.

15. Kolev, N.G., Ullu, E. and Tschudi, C. (2014) The emerging role of RNA-binding proteins in the life cycle of *Trypanosoma brucei*. Cell Microbiol, 16, 482–489.

16. Fernández-Moya, S.M., Carrington, M. and Estévez, A.M. (2014) A short RNA stem–loop is necessary and sufficient for repression of gene expression during early logarithmic phase in trypanosomes. Nucleic Acids Res, gku358.

17. Hendriks, E.F., Robinson, D.R., Hinkins, M. and Matthews, K.R. (2001) A novel CCCH protein which modulates differentiation of *Trypanosoma brucei* to its procyclic form. EMBO J, 20, 6700–6711.

18. Hendriks, E.F. and Matthews, K.R. (2005) Disruption of the developmental programme of *Trypanosoma brucei* by genetic ablation of TbZFP1, a differentiation-enriched CCCH protein. Mol Microbiol, 57, 706–716.

19. Paterou, A., Walrad, P., Craddy, P., Fenn, K. and Matthews, K. (2006) Identification and stage-specific association with the translational apparatus of TbZFP3, a CCCH protein that promotes trypanosome life-cycle development. J Biol Chem, 281, 39002–39013.

20. Walrad, P.B., Capewell, P., Fenn, K. and Matthews, K.R. (2012) The post-transcriptional transacting regulator, TbZFP3, co-ordinates transmission-stage enriched mRNAs in *Trypanosoma brucei*. Nucleic Acids Res, 40, 2869–2883.

21. Benz, C., Mulindwa, J., Ouna, B. and Clayton, C. (2011) The *Trypanosoma brucei* zinc finger protein ZC3H18 is involved in differentiation. Mol Biochem Parasitol, 177, 148–151.

22. Mani, J., Guttinger, A., Schimanski, B., Heller, M., Acosta-Serrano, A., Pescher, P., Spath, G. and Roditi, I. (2011) Alba-domain proteins of *Trypanosoma brucei* are cytoplasmic RNA-binding proteins that interact with the translation machinery. PLoS One, 6, e22463.

23. Subota, I., Rotureau, B., Blisnick, T., Ngwabyt, S., Durand-Dubief, M., Engstler, M. and Bastin, P. (2011) ALBA proteins are stage regulated during trypanosome development in the tsetse fly and participate in differentiation. Mol Biol Cell, 22, 4205–4219.

24. Wurst, M., Seliger, B., Jha, B.A., Klein, C., Queiroz, R. and Clayton, C. (2012) Expression of the RNA recognition motif protein RBP10 promotes a bloodstream-form transcript pattern in *Trypanosoma brucei*. Mol Microbiol, 83, 1048–1063.

25. Kolev, N.G., Ramey-Butler, K., Cross, G.A., Ullu, E. and Tschudi, C. (2012) Developmental progression to infectivity in Trypanosoma brucei triggered by an RNA-binding protein. Science, 338, 1352–1353.

26. Kramer, S., Queiroz, R., Ellis, L., Hoheisel, J.D., Clayton, C. and Carrington, M. (2010) The RNA helicase DHH1 is central to the correct expression of many developmentally regulated mRNAs in trypanosomes. J Cell Sci, 123, 699–711.

27. Gupta, S.K., Kosti, I., Plaut, G., Pivko, A., Tkacz, I.D., Cohen-Chalamish, S., Biswas, D.K., Wachtel, C., Waldman Ben-Asher, H., Carmi, S. et al. (2013) The hnRNP F/H homologue of *Trypanosoma brucei* is differentially expressed in the two life cycle stages of the parasite and regulates splicing and mRNA stability. Nucleic Acids Res, 41, 6577–6594.

28. Vassella, E., Probst, M., Schneider, A., Studer, E., Renggli, C.K. and Roditi, I. (2004) Expression of a major surface protein of *Trypanosoma brucei* insect forms is controlled by the activity of mitochondrial enzymes. Mole Biol Cell, 15, 3986–3993.

29. Hehl, A. and Roditi, I. (1994) The regulation of procyclin expression in *Trypanosoma brucei*: making or breaking the rules? Parasitol Today, 10, 442–445.

30. Hotz, H.R., Hartmann, C., Huober, K., Hug, M. and Clayton, C. (1997) Mechanisms of developmental regulation in *Trypanosoma brucei*: a polypyrimidine tract in the 3′-untranslated region of a surface protein mRNA affects RNA abundance and translation. Nucleic Acids Res, 25, 3017–3026.

31. Mayho, M., Fenn, K., Craddy, P., Crosthwaite, S. and Matthews, K. (2006) Post-transcriptional control of nuclear-encoded cytochrome oxidase subunits in *Trypanosoma brucei*: evidence for genome-wide conservation of life-cycle stage-specific regulatory elements. Nucleic Acids Res, 34, 5312–5324.

32. Archer, S.K., Luu, V.D., de Queiroz, R.A., Brems, S. and Clayton, C. (2009) *Trypanosoma brucei* PUF9 regulates mRNAs for proteins involved in replicative processes over the cell cycle. PLoS Pathog, 5, e1000565.

33. Monk, S.L., Simmonds, P. and Matthews, K.R. (2013) A short bifunctional element operates to positively or negatively regulate ESAG9 expression in different developmental forms of *Trypanosoma brucei*. J Cell Sci, 126, 2294–2304.

34. Wirtz, E., Leal, S., Ochatt, C. and Cross, G.A. (1999) A tightly regulated inducible expression system for conditional gene knock-outs and dominant-negative genetics in Trypanosoma brucei. Mol Biochem Parasitol, 99, 89–101.

35. Hirumi, H. and Hirumi, K. (1989) Continuous cultivation of *Trypanosoma brucei* blood stream forms in a medium containing a low concentration of serum protein without feeder cell layers. J Parasitol, 75, 985–989.

36. Bolger, A.M., Lohse, M. and Usadel, B. (2014) Trimmomatic: a flexible trimmer for Illumina sequence data. Bioinformatics, 30, 2114–2120.

37. Wu, T.D. and Nacu, S. (2010) Fast and SNP-tolerant detection of complex variants and splicing in short reads. Bioinformatics, 26, 873–881.

38. Liao, Y., Smyth, G.K. and Shi, W. (2014) featureCounts: an efficient general purpose program for assigning sequence reads to genomic features. Bioinformatics, 30, 923–930.

39. Robinson, M.D., McCarthy, D.J. and Smyth, G.K. (2010) edgeR: a Bioconductor package for differential expression analysis of digital gene expression data. Bioinformatics, 26, 139–140.

40. McCarthy, D.J., Chen, Y. and Smyth, G.K. (2012) Differential expression analysis of multifactor RNA-Seq experiments with respect to biological variation. Nucleic Acids Res, 40, 4288–4297.

41. Sturn, A., Quackenbush, J. and Trajanoski, Z. (2002) Genesis: cluster analysis of microarray data. Bioinformatics, 18, 207–208.

42. Alexander, D.L., Schwartz, K.J., Balber, A.E. and Bangs, J.D. (2002) Developmentally regulated trafficking of the lysosomal membrane protein p67 in *Trypanosoma brucei*. J Cell Sci, 115, 3253–3263.

43. Kanehisa, M., Araki, M., Goto, S., Hattori, M., Hirakawa, M., Itoh, M., Katayama, T., Kawashima, S., Okuda, S., Tokimatsu, T. et al. (2008) KEGG for linking genomes to life and the environment. Nucleic Acids Res, 36, D480–484.

44. Aslett, M., Aurrecoechea, C., Berriman, M., Brestelli, J., Brunk, B.P., Carrington, M., Depledge, D.P., Fischer, S., Gajria, B. and Gao, X. (2009) TriTrypDB: a functional genomic resource for the Trypanosomatidae. Nucleic Acids Res, 38, D457–D462.

45. Haile, S. and Papadopoulou, B. (2007) Developmental regulation of gene expression in trypanosomatid parasitic protozoa. Curr Opin Microbiol, 10, 569–577.

46. Smith, T.K., Bringaud, F., Nolan, D.P. and Figueiredo, L.M. (2017) Metabolic reprogramming during the *Trypanosoma brucei* life cycle. F1000Res, 6.

47. Reuter, J.S. and Mathews, D.H. (2010) RNAstructure: software for RNA secondary structure prediction and analysis. BMC Bioinformatics, 11, 129.

48. Mulenga, C., Mhlanga, J.D.M., Kristensson, K. and Robertson, B. (2001) *Trypanosoma brucei* brucei crosses the blood-brain barrier while tight junction proteins are preserved in a rat chronic disease model. Neuropath Appl Neuro, 27, 77–85.

49. Voglmayr, J.K., Waites, G. and Setchell, B. (1966) Studies on spermatozoa and fluid collected directly from the testis of the conscious ram. Nature, 210, 861.

50. Cestari, I., Anupama, A. and Stuart, K. (2018) Inositol polyphosphate multikinase regulation of Trypanosoma brucei life stage development. Mol Biol Cell. E17-08-0515.

51. Fenn, K. and Matthews, K.R. (2007) The cell biology of *Trypanosoma brucei* differentiation. Curr Opin Microbiol, 10, 539–546.

52. Silvester, E., McWilliam, K.R. and Matthews, K.R. (2017) The cytological events and molecular control of life cycle development of *Trypanosoma brucei* in the mammalian bloodstream. Pathogens, 6.

53. Fadda, A., Ryten, M., Droll, D., Rojas, F., Farber, V., Haanstra, J.R., Merce, C., Bakker, B.M., Matthews, K. and Clayton, C. (2014) Transcriptome-wide analysis of trypanosome mRNA decay reveals complex degradation kinetics and suggests a role for co-transcriptional degradation in determining mRNA levels. Mol Microbiol, 94, 307–326.

54. Ryley, J. (1962) Studies on the metabolism of the protozoa. 9. Comparative metabolism of blood-stream and culture forms of *Trypanosoma rhodesiense*. Biochem J, 85, 211.

55. Barrett, M.P., Tetaud, E., Seyfang, A., Bringaud, F. and Baltz, T. (1995) Functional expression and characterization of the *Trypanosoma brucei* procyclic glucose transporter, THT2. Biochem J, 312 (Pt 3), 687–691.

56. Morris, M.T., DeBruin, C., Yang, Z.Q., Chambers, J.W., Smith, K.S. and Morris, J.C. (2006) Activity of a second *Trypanosoma brucei* hexokinase is controlled by an 18-amino-acid C-terminal tail. Eukaryot Cell, 5, 2014–2023.

57. Bakker, B.M., Walsh, M.C., ter Kuile, B.H., Mensonides, F.I., Michels, P.A., Opperdoes, F.R. and Westerhoff, H.V. (1999) Contribution of glucose transport to the control of the glycolytic flux in *Trypanosoma brucei*. Proc Natl Acad Sci U S A, 96, 10098–10103.

58. Ashe, M.P., De Long, S.K. and Sachs, A.B. (2000) Glucose depletion rapidly inhibits translation initiation in yeast. Mol Biol Cell, 11, 833–848.

59. Coller, J. and Parker, R. (2005) General translational repression by activators of mRNA decapping. Cell, 122, 875–886.

60. Teixeira, M.C., Duque, P. and Sa-Correia, I. (2007) Environmental genomics: mechanistic insights into toxicity of and resistance to the herbicide 2,4-D. Trends Biotechnol, 25, 363–370.

61. Gancedo, J.M. (2008) The early steps of glucose signalling in yeast. FEMS Microbiol Rev, 32, 673–704.

62. Kuhn, K.M., DeRisi, J.L., Brown, P.O. and Sarnow, P. (2001) Global and specific translational regulation in the genomic response of *Saccharomyces cerevisiae* to a rapid transfer from a fermentable to a nonfermentable carbon source. Mol Cell Biol, 21, 916–927.

63. Andreev, D.E., O’Connor, P.B., Zhdanov, A.V., Dmitriev, R.I., Shatsky, I.N., Papkovsky, D.B. and Baranov, P.V. (2015) Oxygen and glucose deprivation induces widespread alterations in mRNA translation within 20 minutes. Genome Biol, 16, 90.

64. Priest, J.W. and Hajduk, S.L. (1994) Developmental regulation of mitochondrial biogenesis in *Trypanosoma brucei*. J Bioenerg Biomembr, 26, 179–191.

65. Kligun, E. and Mandel-Gutfreund, Y. (2015) The role of RNA conformation in RNA-protein recognition. RNA Biol, 12, 720–727.

66. Allers, J. and Shamoo, Y. (2001) Structure-based analysis of Protein-RNA interactions using the program ENTANGLE. J Mol Biol, 311, 75–86.

67. de Smit, M.H. and van Duin, J. (1990) Secondary structure of the ribosome binding site determines translational efficiency: a quantitative analysis. Proc Natl Acad Sci U S A, 87, 7668–7672.

68. Wen, J.D., Lancaster, L., Hodges, C., Zeri, A.C., Yoshimura, S.H., Noller, H.F., Bustamante, C. and Tinoco, I. (2008) Following translation by single ribosomes one codon at a time. Nature, 452, 598–603.

69. Qu, X., Wen, J.D., Lancaster, L., Noller, H.F., Bustamante, C. and Tinoco, I., Jr. (2011) The ribosome uses two active mechanisms to unwind messenger RNA during translation. Nature, 475, 118–121.

70. Jagodnik, J., Chiaruttini, C. and Guillier, M. (2017) Stem-loop structures within mRNA coding sequences activate translation initiation and mediate control by small regulatory RNAs. Mol Cell, 68, 158–170 e153.

